# A spatial code in the dorsal lateral geniculate nucleus

**DOI:** 10.1101/473520

**Authors:** Vincent Hok, Pierre-Yves Jacob, Pierrick Bordiga, Bruno Truchet, Bruno Poucet, Etienne Save

**Affiliations:** Aix Marseille Univ, CNRS, LNC, Marseille, France

**Keywords:** Hippocampus, dorso-Lateral Geniculate Nucleus, Vision, Spatial Cognition, Place Cells

## Abstract

Since their discovery in the early ‘70s^1^, hippocampal place cells have been studied in numerous animal and human spatial memory paradigms^2–4^. These pyramidal cells, along with other spatially tuned types of neurons (e.g. grid cells, head direction cells), are thought to provide the mammalian brain a unique spatial signature characterizing a specific environment, and thereby a memory trace of the subject’s place^5^. While grid and head direction cells are found in various brain regions, only few hippocampal-related structures showing ‘place cell’-like neurons have been identified^6,7^, thus reinforcing the central role of the hippocampus in spatial memory. Concurrently, it is increasingly suggested that visual areas play an important role in spatial cognition as recent studies showed a clear spatial selectivity of visual cortical (V1) neurons in freely moving rodents^8–10^. We therefore thought to investigate, in the rat, such spatial correlates in a thalamic structure located one synapse upstream of V1, the dorsal Lateral Geniculate Nucleus (dLGN), and discovered that a substantial proportion (ca. 30%) of neurons exhibits spatio-selective activity. We found that dLGN place cells maintain their spatial selectivity in the absence of visual inputs, presumably relying on odor and locomotor inputs. We also found that dLGN place cells maintain their place selectivity across sessions in a familiar environment and that contextual modifications yield separated representations. Our results show that dLGN place cells are likely to participate in spatial cognition processes, creating as early as the thalamic stage a comprehensive representation of one given environment.

---

We recorded neurons from 9 rats trained in a pellet-chasing task in a circular arena. The animals were either implanted above the CA3 region (in order to record CA3 place cells and then dLGN neurons; *n* = 5) or just above the dLGN (*n* = 4) in order to further characterize dLGN place cell properties. Overall, we recorded 228 CA3 and 196 dLGN place cells (see Fig. 1a—e for recording locations and Extended Data Tab. 1 for detailed results per animal).

**Figure 1.**
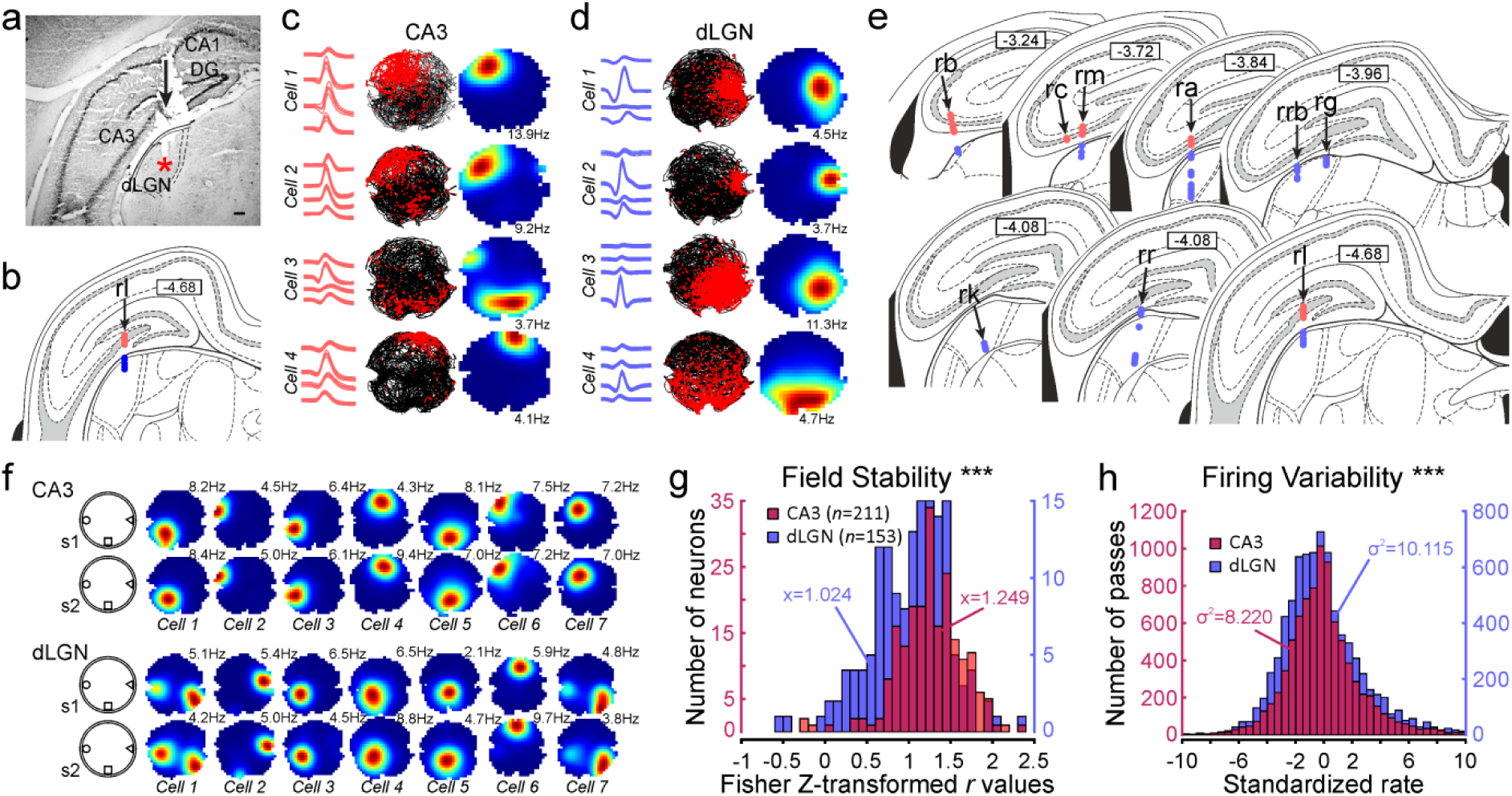
Recording locations and basic spatial properties of CA3 and dLGN place cells. ***a,*** Micrograph showing the trajectory of the electrodes bundle (arrow) and the final position of the tip of the bundle (red asterisk). Scale bar = 200µm. CA1, CA2 and CA3: Cornu Ammonis subfields; DG: Dentate Gyrus; dLGN: Dorsal Lateral Geniculate nucleus. ***b,*** Corresponding atlas section indicating the location of CA3 place cells (red circles) and dLGN place cells (blue circles) recordings. ***c,*** CA3 place cell examples showing waveforms (left column), trajectory maps (middle) and rate maps (right). Rate maps are coded according to a color scale that goes from blue (silent) to red (maximum rate), with cyan, green, yellow, and orange pixels as intermediate firing rates from low to high. Numbers located on the right side of the maps indicate average activity within the firing field. ***d,*** dLGN place cell examples. Note the typical pyramidal waveform shape observed for CA3 place cells recordings whereas dLGN waveforms show narrower, bigger and usually biphasic waveforms. ***e,*** Coronal sections of the rat brain showing the location of recorded place cells in the hippocampus (red circles) and the dLGN (blue circles) for all animals. Arrows indicate the trajectory of the electrodes bundle for each rat (rats are identified with 2 or 3 letters). For rats where CA3 and dLGN place cells were recorded successively (*n* = 4), the average minimal distance between the deepest CA3 place cell and the most superficial dLGN place cell recorded was 246.09 ± 63.28 µm, which is compatible with the anatomical distance between the two structures. Numbers in inserts indicate the stereotaxic coordinates following the anterior-posterior axis. ***f,*** Examples of CA3 and dLGN place cells (each set of cells extracted from three different rats), both showing good stability across successive sessions. ***g,*** Histogram showing Fisher Z-transformed correlation values between successive sessions for dLGn (blue) and CA3 (red) place cells. Although dLGN place fields appear less stable than CA3, both populations show strong stability across time (*** *p*<0.001). ***h,*** dLGN place cells show greater firing variability (overdispersion) than CA3 units (*** *p*<0.001).

We first confirmed our histology-based identification with an analysis of electrophysiological properties extracted from neurons waveforms (Extended Data Fig. 1a). To this end we used the coefficient of variation and the peak-to-trough parameters that allowed us to isolate two main clusters (Extended Data Fig. 1b). Using k-means clustering, we identified a first cluster that comprised around 75% of the CA3 place cells and a second cluster that comprised around 93% of dLGN place cells (insert in Extended Data Fig. 1b). To rule out the possibility of fibers origin to our dLGN recordings we looked at the types of waveforms recorded in this region. We therefore computed a normalized phase plot (which shows the rate of change of the membrane potential (*dV/dt*) against the membrane potential itself) for each waveform and computed the barycentre of the resulting polygon (Extended Data Fig. 1c *top*). We then reported the *x* and *y* coordinates of the barycentre on a two-dimensional plot (Extended Data Fig. 1c *bottom*). According to this representation, triphasic waveforms, usually considered as compatible with action potentials emitted by fibers^11^, should have their barycentre coordinates clustered around the plot origin (Extended Data Fig. 1d). Here, we found scarce recordings displaying such triphasic waveforms (Extended Data Fig. 1e), thus excluding the fibers origin of our recordings. Using k-means clustering, we identified two clusters based on dLGN data. A first cluster was constituted mainly from biphasic waveforms (negative-positive potential) and the second from mainly monophasic (positive potential) waveforms (insert in Extended Data Fig. 1e). As no difference was observed regarding the spatial characteristics between these two groups of neurons (data not shown), we pooled the data into a single dLGN population. Therefore, we considered hereafter only two populations of place cells (CA3 *vs.* dLGN) for simple comparisons on basic place cell properties (summarized in Table 1). dLGN place cells mainly differed from CA3 place cells according to their overall firing activity (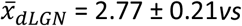. 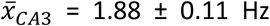, two-sample t-test assuming unequal variances, *t*_(296)_ = −3.8474, *p* = 1.4625.10^-4^), a difference that might be explained by dLGN greater primary field size (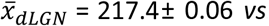. 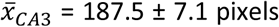 pixels, *t*_(340)_ = −2.2886, *p* = 0.0227) and number of place fields (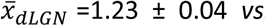. 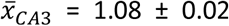 place fields, *t*_(308)_ = −3.6608, *p* = 2.9578.10^-4^). Given these differences, information content logically was markedly decreased in dLGN place cells (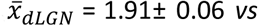. 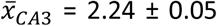 bits per spike, *t*_(409)_ = 4.7384, *p* = 2.9772.10^-6^). To some extent, the greater number of dLGN place fields appeared related to the presence of objects placed at the periphery of the arena. Removing the objects while maintaining a cue card attached to the surrounding curtains did not affect overall place field stability (Extended Data Fig. 2a—c), but some place fields in the vicinity of objects disappeared (see for instance cells #4 and #7 in Extended Data Fig. 2a and population comparisons in Extended Data Tab. 2). This last result raises the possibility that some dLGN cells act as object cells similar to what has been observed in other brain structures^6,12^.

**Table 1.** Main electrophysiological and spatial characteristics of CA3 and dLGN place cells.

|  | CA3<br>(n=228) | dLGN<br>(n=196) |
| --- | --- | --- |
| **Waveform Characteristics** |  |  |
| Spike Amplitude (μV) | 215.8 ± 9.5 | 375.6 ± 19.7*** |
| Spike Height (μV) | 156.4 ± 9.5 | 342.9 ± 18.7*** |
| Spike Half-width (μs) | 279.4 ± 5.5 | 179.7 ± 6.1*** |
| Peak-to-trough (μs) | 357.2 ± 8.9 | 150.3 ± 6.0*** |
| **Firing Rate (Hz)** |  |  |
| Overall | 1.88 ± 0.11 | 2.77 ± 0.21*** |
| Infield (GR) | 6.64 ± 0.26 | 6.55 ± 0.26 |
| Infield (CR) | 21.3 ± 1.00 | 22.6 ± 1.15 |
| **Spatial Characteristics** |  |  |
| Spatial Coherence | 0.75 ± 0.01 | 0.75 ± 0.01 |
| IC (bits/spike) | 2.24 ± 0.05 | 1.91 ± 0.06*** |
| Number of fields | 1.08 ± 0.02 | 1.23 ± 0.04*** |
| Field Size (pixels) | 187.5 ± 7.1 | 217.4 ± 11.1* |

Spike Height: Peak relative to resting potential.
Spike Amplitude: Peak relative to the most negative voltage reached during afterhyperpolarization immediately after the spike.
Spike Half-width: Width at half-maximal spike height.
GR: Grand Rate; CR: Center Rate; IC: Information Content.
Averages are given ± SEM. \*\*\*, $p < 0.001$ ; \*, $p < 0.05$

For cells recorded during multiple sessions on a single day, we analysed field stability for both dLGN (*n* = 153) and CA3 (*n* = 211) place cells (Fig. 1f). Both populations showed remarkable field stability between successive recording sessions (Fig. 1g), although CA3 place field stability remained significantly higher than dLGN (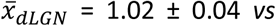. 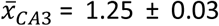, *t*_(295)_ = −4.8582, *p* = 1.9266.10^-6^). Since hippocampal place cell firing variability (*i.e.* overdispersion) has been suggested to reflect an internally generated cognitive process^13–15^ we looked at this phenomenon in our dLGN recordings. We found a much greater overdispersion in dLGN place cells (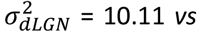. 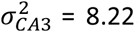, Levene’s test for equality of variances, W = 59.6356, *p* = 1.2101.10^-14^, Fig. 1h) suggesting that dLGN place cells are sensitive to extra-positional information much like hippocampal place cells.

In the next step, we asked whether phase precession, a process usually interpreted as the prediction of the sequence of upcoming positions, could be observed in dLGN place cells. Because such process relies heavily on the presence of theta oscillations (4-12 Hz), we first analysed their characteristics in the local field potentials (LFPs) obtained in both CA3 and dLGN. The dLGN LFPs showed a much higher theta peak frequency than CA3 (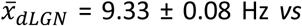. 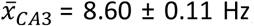, *t*_(129)_ = - 5.3904, *p* = 3.2325.10^-7^) while the relative power in the theta band was not different (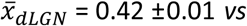. 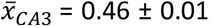, two-sample t-test, *t*_(151)_ = 1.7755, *p* = 0.0778; Fig. 2a—c). Intrinsic firing peak in the theta band was lower in dLGN place cells (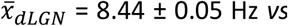Hz *vs.* 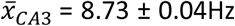Hz, *t*_(315)_ = 4.5412, *p* = 7.9734.10^-6^; Fig. 2d and 2e). These results suggest that phase precession is less likely to occur in dLGN. This was confirmed by comparing the proportions of phase precessing place cells in both structures (%_*dLGN*_ = 10.79 *vs.* %_*CA3*_ = 27.53, 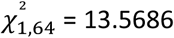, *p* = 2.2995.10^-4^; Fig. 2f and 2g). Comparing the circular-linear correlation coefficients (*rho* values) revealed a significant difference, the relation between phase and position of spikes was weaker in dLGN (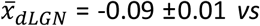. 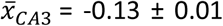, two-sample t-test, *t*_(62)_ = −2.3605, *p* = 0.0214; Fig. 2h). These results indicate that one of the key features of spatial predictive coding is virtually absent from dLGN place cell activity, thus suggesting a contribution to spatial information processing functionally distinct from CA3.

**Figure 2.**
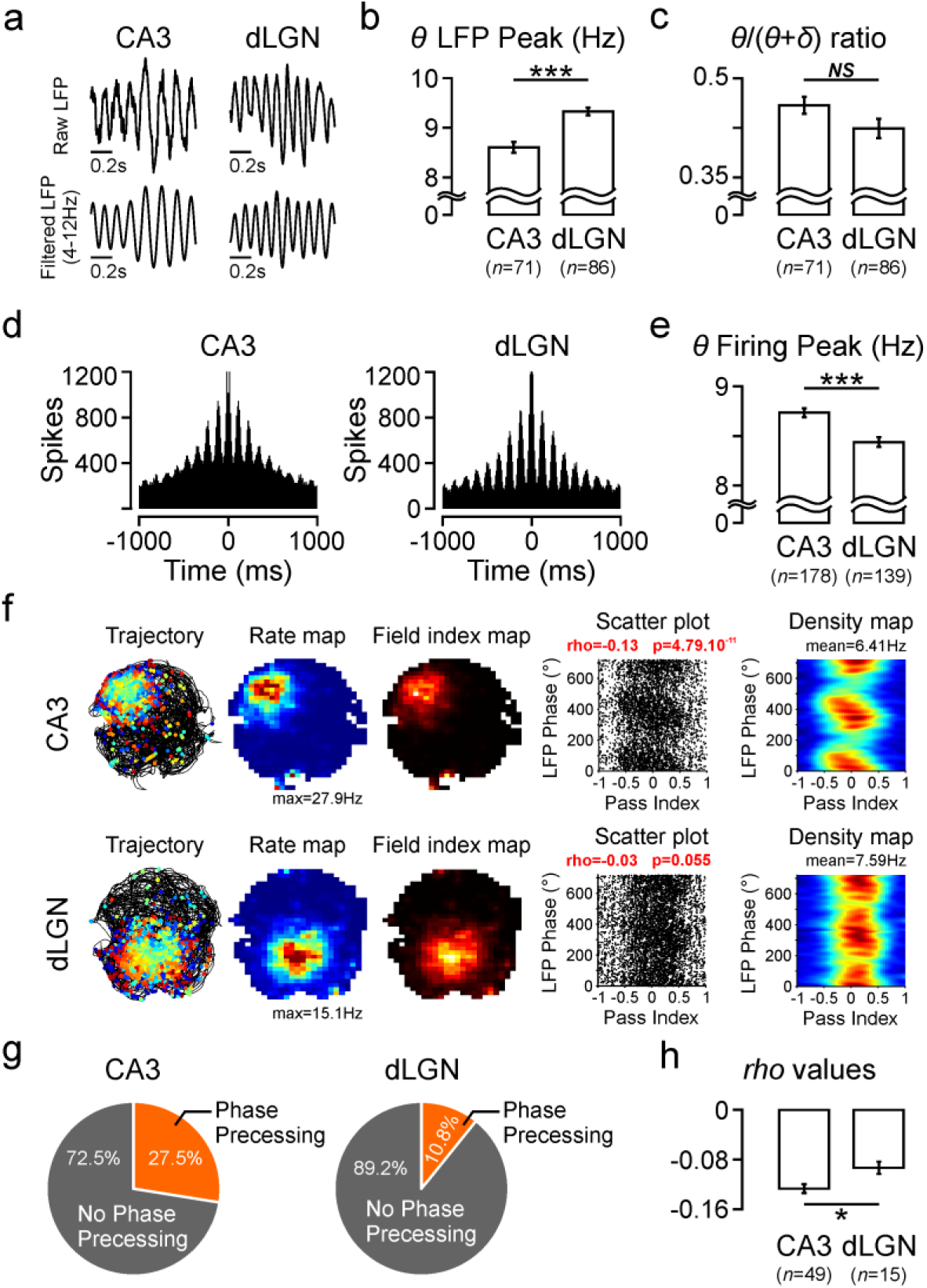
Comparison of spike-theta phase relationships in CA3 and dLGN place cells. ***a,*** Examples of raw (top row) and filtered (4-12Hz, bottom row) LFP recorded in CA3 and dLGN. ***b,*** Average peak frequency (± SEM) in the theta band (4-12Hz) of LFPs for both structures (*** *p*<0.001). ***c,*** Average Theta/(Theta+Delta) ratio in CA3 and dLGN recordings (*n.s.*). ***d,*** Autocorrelograms of the firing activity from one CA3 place cell and one dLGN place cell recorded along with the local field potentials shown in ***a***. ***e,*** Average peak frequency in the theta band of the place cell autocorrelograms for both structures (*** *p*<0.001). ***f,*** Example plots from one precessing CA3 place cell (top row; linear–circular correlation, *p*<0.01) and from one dLGN non-precessing place cell (bottom row). From *left* to *right*: Trajectory with spikes colored by the pass index at each spike, rate map, field index map, scatter plot of the LFP theta phase *vs.* the pass index and occupancy-normalized rate map of the scatter plot (see Supplementary Methods). Correlation coefficients (rho values) and significance (*p* values) are indicated in red. ***g,*** Proportion of phase precessing cells in CA3 and dLGN. ***h,*** Average circular-linear correlation coefficient (rho values) for phase precessing cells identified in CA3 and dLGN.

In a third step, we tested in two additional animals whether dLGN cells would respond to a simple visual stimulation, hereby indicating that the electrodes were indeed located in a structure that processes visual information. In this experiment, we recorded dLGN cells in both normal and 1 Hz flickering lighting conditions (Fig. 3a). We obtained a total of 64 dLGN cells in two animals over 13 sessions where place cells and 1Hz modulated cells were recorded simultaneously (Fig. 3b and 3c and Extended Data Fig. 2d). Out of these 64 dLGN cells, we identified 25% of units significantly modulated at 1Hz, and nearly 30% of place cells (Fig. 3d). In order to compare the spatial selectivity between these two populations, we computed the spatial sparsity (see Supplementary Methods) for each cell. In this case, the sparsity value ranges between zero (for a uniform rate map) and 1 (for a delta function rate map)^16^. This analysis allowed us to confirm that none of the 1Hz modulated cells could be identified as a spatial selective cell (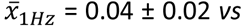. 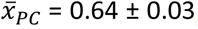, two-sample t-test, *t*_(27)_ = 17.9833, *p* = 1.4798.10^-16^; Fig. 3e). Therefore, dLGN place cells were insensitive to this particular visual stimulation but were nonetheless recorded simultaneously with an important proportion of cells that did respond.

**Figure 3.**
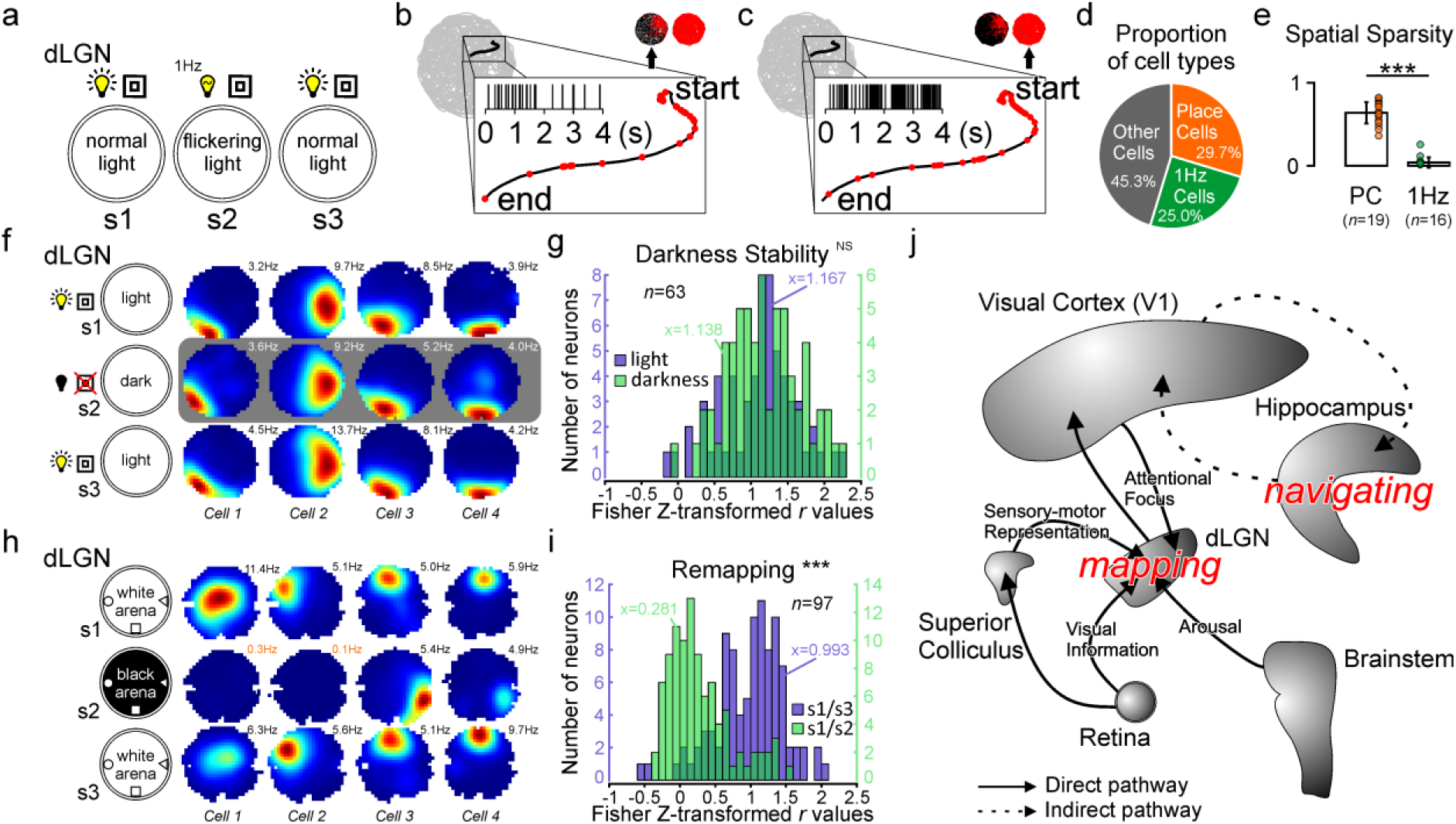
Further characterization of dLGN place cells properties. ***a,*** Schematic showing the procedure used to assess visual entrainment of dLGN units in freely moving animals. dLGN cells were recorded during normal lighting conditions (s1 and s3) and during exposure to a flickering ambient light oscillating at 1Hz (s2). ***b,*** Close up of one pass (insert) for a dLGN place cell recorded during the flickering light session. The raster plot obtained for the duration of the pass does not show any clear firing modulation at 1Hz. ***c,*** Same as previously for a dLGN cell recorded simultaneously with place cell recorded in ***b***. This time, the raster plot shows a strong modulation at 1Hz. ***d,*** Proportion of cell types recorded in the experiment presented previously. Note that only sessions with dLGN place cells and 1Hz modulated cells recorded simultaneously were kept for this analysis. ***e,*** Comparison of spatial sparsity between place cells and 1Hz modulated cells. This bar graph shows that none of the identified place cells are entrained by visual stimulation and that none of the identified 1Hz modulated cells are place cells (*** *p*<0.001). ***f,*** Subset of dLGN place cells recorded in light-dark-light conditions. The animals were left in the arena and no odor removal procedure was performed between successive sessions. A vast majority of dLGN place cells maintained their spatial firing activity in complete darkness. ***g,*** Histogram showing Fisher Z-transformed correlations values between light-light (blue bars) and light-darkness (green bars) conditions (*NS*). ***h,*** Subset of dLGN place cells recorded in two environments differing in color settings (wall and floor). Some cells cease firing during the second session (*e.g.* cells 1 and 2; orange numbers indicate the overall average firing rate of the cell when no place field is detected) while others change their firing location and rate activity (*e.g.* cells *3* and 4). ***i,*** Histogram showing Fisher Z-transformed correlations values between similar (blue bars) and different (green bars) environments (*** *p*<0.001). ***j,*** Schematic showing the principal structures involved in visual and spatial processing in the rat brain. The dLGN appears fully qualified to elaborate the mapping of the environment through its connections with the retina (visual), superior colliculus (sensory-motor), brainstem (arousal) and primary visual cortex (attention). According to this view, the primary role of the hippocampus would be devoted to the navigational component of a global navigation system, elaborating possible routes based on dLGN mapping information. See supplementary discussion for further considerations on this specific matter.

Considering the primary role of dLGN in visual processing, we then asked if dLGN place cell spatial selectivity was maintained in the absence of visual information. Therefore, we recorded a subset of dLGN place cells (*n* = 63) in a circular environment with a distal cue card attached to the surrounding curtain in normal lighting conditions and in total darkness (Fig. 3f). In between sessions, the animals were left in the arena and no attempt was made to neutralize odor cues. The vast majority (*ca.* 90%) of dLGN place cells showed stable spatial activity in both light-light and light-darkness conditions (Fisher Z-transformed correlation values, 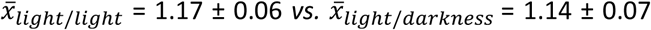, paired t-test, *t*_(62)_ = 0.6108, *p* = 0.5435, *NS*; Fig. 3g). This result demonstrates that dLGN spatially localized firing patterns, once learned, are not likely to depend upon local visual cues within the experimental room.

In order to assess the control of objects exerted on dLGN spatial representation, we recorded a small subset of place cells in standard and rotated (90° clockwise rotation of object set while the animal was out of the arena) conditions. All the dLGN cells tested in this experiment followed the rotation with their place fields rotated to an equal amount (Extended Data Fig. 2e—g).

Given the striking similarities between dLGN and CA3 place cells with respect to their spatial characteristics, we further asked whether dLGN place cells would show more complex features classically reported in CA3. Hippocampal place cells are well known to change their spatial firing activity when the animal is introduced in distinct environments differing either in their shape, color or both^17,18^. This phenomenon, known as global remapping, is thought to reflect mechanisms involved in decorrelation of overlapping inputs^19^. Such processes were observed in dLGN place cells (Fig. 3h). Some dLGN place cells ceased firing while the animals were introduced in an alternate environment (see for example cells #1 and 2 in Fig. 3h) or changed their firing location (cells #3 and 4), as previously observed in hippocampal place cells^17,18^. Overall, the population of dLGN place cells showed great place field stability between similar environments and complete remapping between distinct environments (Fisher Z-transformed correlation values, 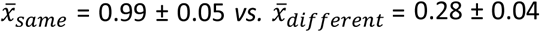, paired t-test, *t*_(96)_ = −12.5846, *p* = 4.9708.10^-22^; Fig. 3i) similar to what has been found in hippocampal place cells^13^.

Our results show that most of the prerequisites for spatial mapping (e.g. spatial selectivity, stability, environment specificity) are present at the dLGN level. Visual and navigational systems are known to closely interact in both rodent^8,9^ and primate brain^20,21^, but the boundaries between these two systems are becoming less and less clearly delimited^10,22^. Additionally, when looking at the great diversity of dLGN inputs^23^, one can surmise that this thalamic structure is not solely dedicated to visual processing (Fig. 3j, see also Supplementary Discussion). Indeed, the nature of converging cortical^24–26^ and subcortical^27–29^ inputs on dLGN is likely giving birth to the true original spatial mapping function of the mammalian brain.

## Methods

Many of the procedures used here are described elsewhere. For further details, the reader is referred to Hok et al.^13^. All procedures complied with both European (directive 2010/63/EU of the European Parliament and of the Council) and French (AGRG1238767A) institutional guidelines.

### Subjects

Male Long-Evans rats (Centre d’Élevage Janvier, St.-Berthevin, France), weighing 300-350 g, were housed in separate cages at 20° C under a 12 hr light/dark cycle. After being handled for two weeks, animals were food-deprived to 85% of their bodyweight and trained in the pellet chasing task.

### Apparatus and Behavioral Procedures

The animals were trained in two cylindrical environments that differed in color (floor and wall) with three different objects that differed in color, shape and texture, located at the periphery of the apparatus and disposed in isosceles triangle configuration. This particular configuration of object is known to efficiently control the firing fields of hippocampal place cells^30^. Four copies of each object were available so that a different set of objects could be used on each session to avoid association of olfactory cues with a specific environment. The recording arenas had a 76 cm diameter with a 50 cm high wall, were surrounded by ceiling-high curtains and were dimly lit from above (luminance was kept constant at 10 lux). During screening phases, the animal was placed in a flower pot (22 cm in diameter with a 40 cm high wall) located close by the arena. A wireless speaker placed at the vertical of the arena was continuously diffusing a musical background in order to mask uncontrolled auditory cues. Unless otherwise stated, the floor was wiped clean with a damp cloth imbibed with alcohol in between two sessions in order to neutralize olfactory cues left on the floor of the apparatuses.

Subjects were then placed in the open field and 20 mg food pellets (TestDiet™, 5TUL formula) were distributed every 20 seconds to random locations within the open field by an automatic pellet dispenser (ENV-203M, Med Associates Inc.); in this way, animals were in continuous locomotion, allowing for complete sampling of the environment. Each recording session lasted 20 minutes.

### Electrodes

Recording electrodes were assembled in a bundle of eight tetrodes platinum-iridium wires (90% platinum, 10% iridium; HML insulated, 25μm bare wire diameter, California Fine Wire Ltd., California, USA). Tetrodes were threaded through a guide 21 gauge cannula. Tetrodes were then mounted on a lightweight microdrive, cut flat and implanted either in the dorsal hippocampus, above the CA3 subfield (−4.0 AP, ±4.0 ML and 3.0 mm dorsoventral to dura; 5 rats) or just above dLGN (−4.0 AP, ±4.0 ML and 3.4 mm dorsoventral to dura; 4 rats). The microdrives used here are built around three precision screws, machined to a pitch of 450 µm, which advance the electrodes in 30 µm steps (1/16th turns). The electrode headstage was then anchored to the skull using miniature titanium screws embedded in dental resin cement.

## Extended Data

**Extended Data Figure 1.**
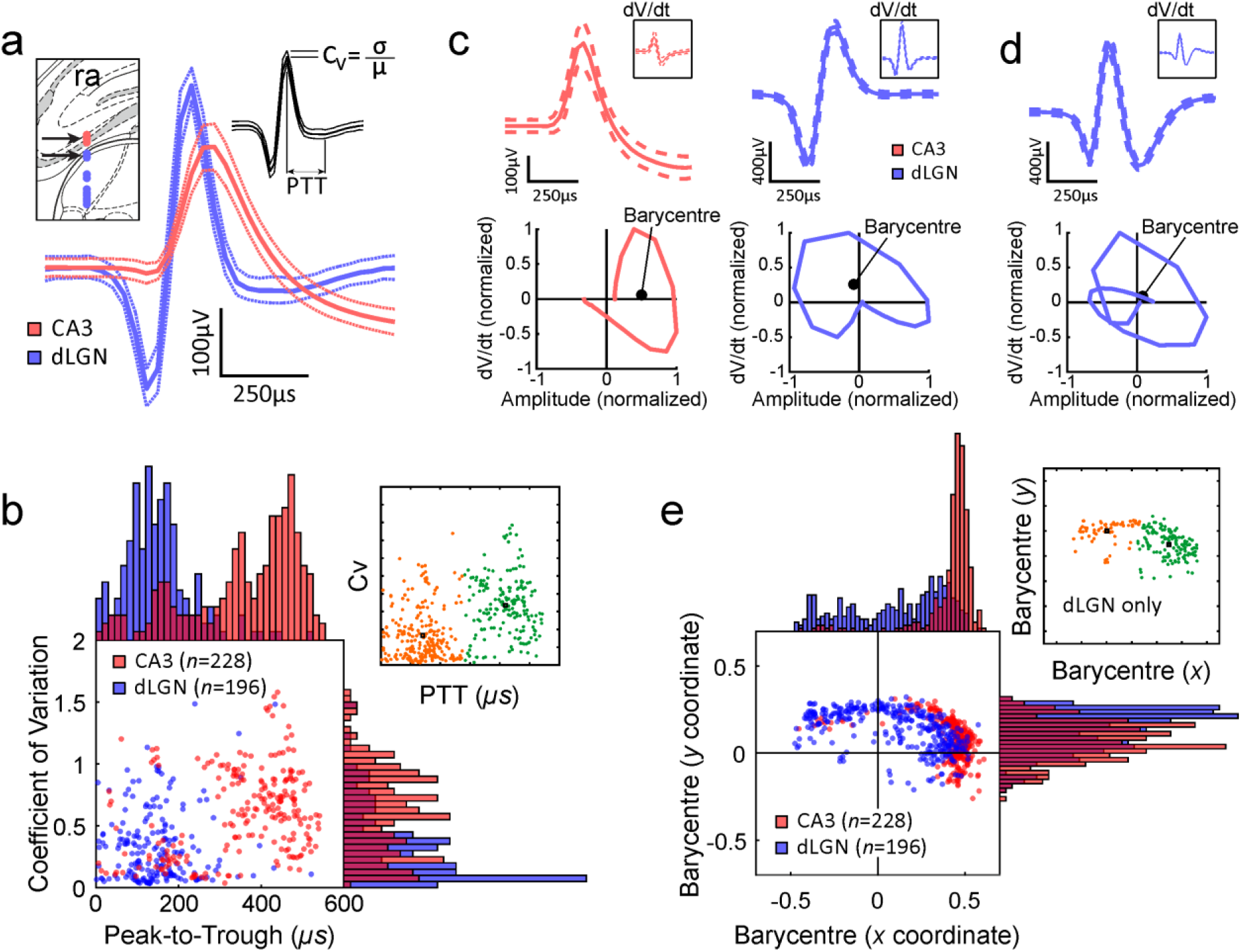
Waveform properties of CA3 and dLGN place cells. ***a,*** Representative waveforms of hippocampal (red trace) and dLGN place cells (blue trace) recorded in the same animal (location indicated by arrows in insert). For each waveform, the solid line is the average waveform shape, and the dashed lines show the 1 SD confidence intervals. ***b,*** Clear dissociation of the two populations of cells based on their waveform characteristics. The scatter plot is showing the distribution of the hippocampal (red) and dLGN (blue) place cell populations that are easily segregated based on their peak-to-trough time (PTT) and coefficient of variation (C_V_). ***c,*** Construction of normalized phase plots for one CA3 (red traces) and one dLGN (blue traces) place cells. For each waveform (top) the *x* and *y* coordinates of the polygon barycentre are computed (bottom). ***d***, Example of one dLGN triphasic waveform shape (top) and the resulting phase-plot (bottom). The barycentre coordinates are located around the plot origin. Note however the high amplitude of the signal, hardly compatible with axonal action potential origin. ***e,*** The scatter plot is showing the distribution of the hippocampal (red) and dLGN (blue) barycentres coordinates. This two-dimensional visualization shows the wide diversity of dLGN waveforms compared to CA3.

**Extended Data Figure 2.**
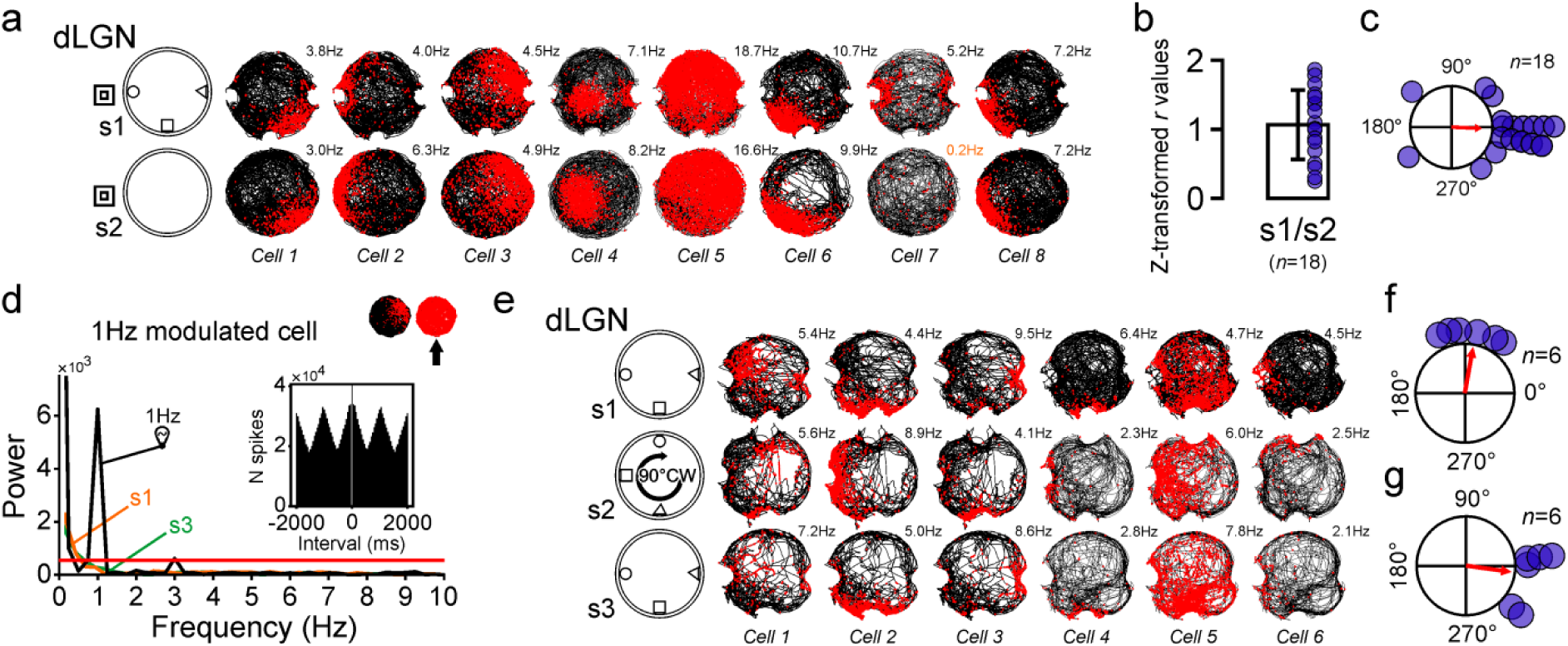
dLGN place cells responses to environmental manipulations. ***a,*** Examples of dLGN place cells recorded in object and cue conditions (first session, s1) and after object removal (second session, s2). The animals were removed from the arena in between the two sessions and the floor wiped clean with a damp cloth imbibed with alcohol. Note that the vast majority of place fields are maintained after object removal and only few cells cease firing at former object location (*e.g.* cells 4 and 7). ***b,*** Bar graph showing the distribution and average of the Fisher Z-transformed correlation values obtained when comparing the first and second sessions. Overall, the 18 place cells tested in these conditions showed a good spatial stability. ***c,*** Polar plot showing the distribution of the angles at which the correlations between the two sessions were maximal. A *V* test for circular uniformity confirmed that the angle values are clustered around 0° (*v* = 13.2944, *p* = 4.6797.10^-6^). The red arrow indicates the vector length with a value of 0.7392 and a mean direction of −2.3768°. Overall, removing the objects did not affect place field stability. ***d,*** Example of a fast fourier transform based on autocorrelogram (insert) showing a strong 1Hz modulation during visual entrainment. The red line indicates the significance threshold. Same cell as presented in Fig. 2c. ***e,*** Six dLGN place cells were recorded in standard conditions (first and third sessions) and following 90° clockwise rotation of the set of objects (second session). The animals were removed from the arena between two sessions and the floor wiped clean with a damp cloth imbibed with alcohol. ***f,*** Polar plot showing the distribution of the angles at which the correlations between the two first sessions (s1 and s2) were maximal. The *V* test for circular uniformity indicated that the angle values are clustered around 90° (*v* = 4.5414, *p* = 0.0044). The red arrow indicates the vector length with a value of 0.9450 and a mean direction of 79.8417°. ***g,*** Polar plot showing the distribution of the angles at which the correlations between the two standard sessions (s1 and s3) were maximal. The *V* test confirmed that the angle values are clustered around 0° (*v* = 5.4865, *p* = 7.6148.10^-4^; vector length = 0.9233, mean direction = −7.9340°).

**Extended Data Table 1.** Total number of sessions and cells recorded in CA3 and dLGN per subject.

| **Subject** | **#Sessions** |  | **#Total Cells** |  | **#Place Cells** |  | **%Place Cells** |  |
| --- | --- | --- | --- | --- | --- | --- | --- | --- |
|  | **CA3** | **dLGN** | **CA3** | **dLGN** | **CA3** | **dLGN** | **CA3** | **dLGN** |
| ra | 7 | 21 | 51 | 104 | 11 | 26 | 21.6% | 25.0% |
| rb | 18 | 3 | 243 | 13 | 57 | 4 | 23.5% | 30.8% |
| rl | 34 | 5 | 295 | 15 | 75 | 7 | 25.4% | 46.7% |
| rm | 25 | 6 | 251 | 12 | 83 | 7 | 33.1% | 58.3% |
| rc | 2 |  | 21 |  | 2 |  | 9.5% |  |
| rg |  | 16 |  | 89 |  | 37 |  | 41.6% |
| rk |  | 23 |  | 108 |  | 32 |  | 29.6% |
| rr |  | 34 |  | 223 |  | 59 |  | 26.5% |
| rrb |  | 12 |  | 52 |  | 24 |  | 46.1% |
| **Total** | **86** | **120** | **861** | **616** | **228** | **196** | **26.5%** | **31.8%** |

**Extended Data Table 2.**
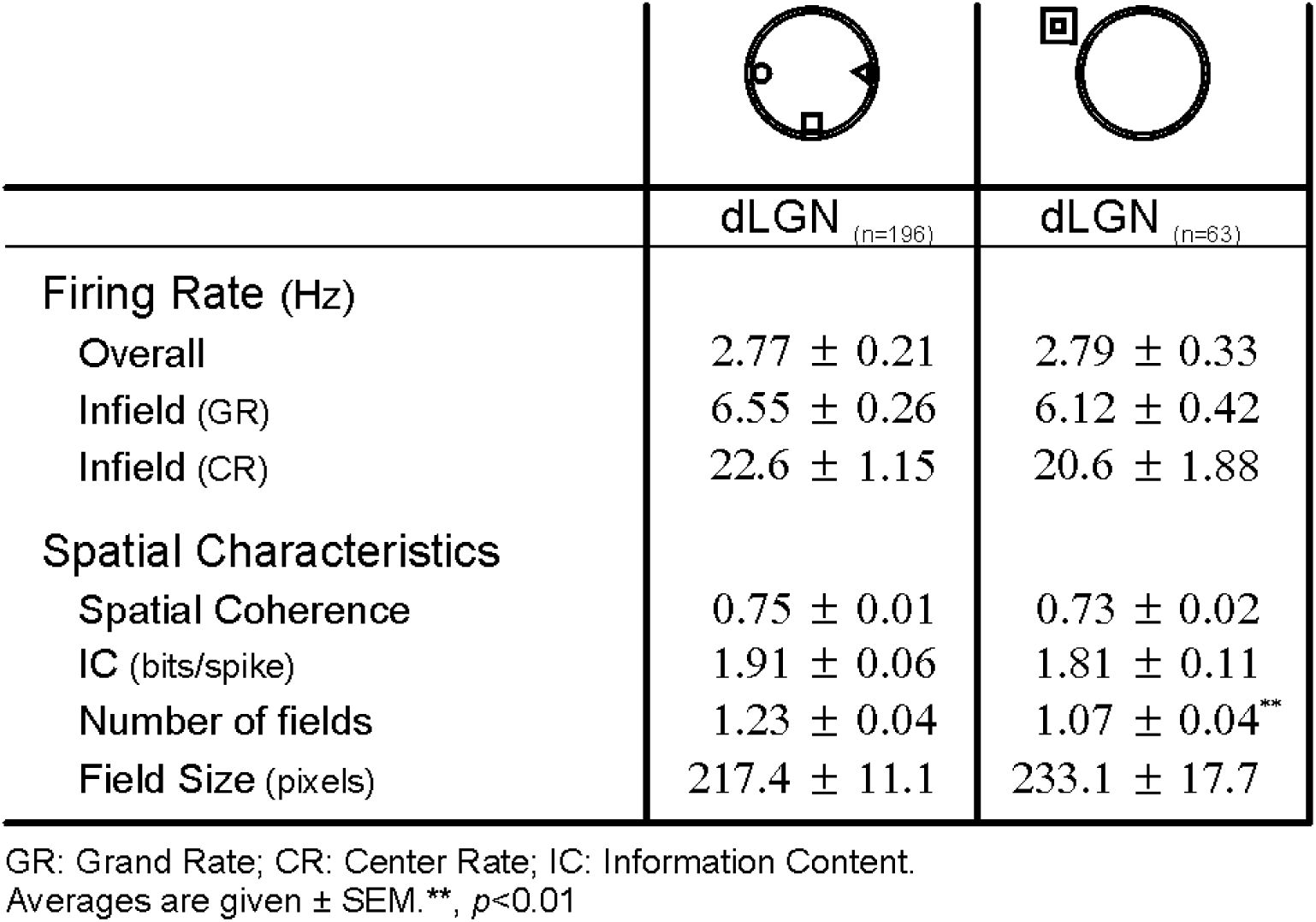
Main electrophysiological and spatial characteristics of dLGN place cells with or without objects.

|  |  |  |
| --- | --- | --- |
|  | dLGN (n=196) | dLGN (n=63) |
| **Firing Rate (Hz)** |  |  |
| Overall | 2.77 ± 0.21 | 2.79 ± 0.33 |
| Infield (GR) | 6.55 ± 0.26 | 6.12 ± 0.42 |
| Infield (CR) | 22.6 ± 1.15 | 20.6 ± 1.88 |
| **Spatial Characteristics** |  |  |
| Spatial Coherence | 0.75 ± 0.01 | 0.73 ± 0.02 |
| IC (bits/spike) | 1.91 ± 0.06 | 1.81 ± 0.11 |
| Number of fields | 1.23 ± 0.04 | 1.07 ± 0.04** |
| Field Size (pixels) | 217.4 ± 11.1 | 233.1 ± 17.7 |

GR: Grand Rate; CR: Center Rate; IC: Information Content. Averages are given ± SEM. \*\*, *p*<0.01

## Supplementary Methods

### Surgical procedure

Rats were deeply anesthetized by an intra-peritoneal injection of a mixture of an α2 adrenergic agonist (medetomidine, Domitor^®^, 0.25 mg/kg) and a NMDA receptor antagonist (ketamine, Imalgene^®^, 60 mg/kg). Following surgery, all animals received subcutaneous injections of a broad-spectrum antibiotic (oxytetracycline, Terramycine^®^, 60 mg/kg) and a nonsteroidal anti-inflammatory drug (tolfenamic acid, Tolfedine^®^, 4 mg/kg) and were allowed one week to recover.

### Recording Procedures

Initial amplification of extracellular voltage was performed by a headstage connected to the electrode housing. A cable carried the signals to a powered slip ring commutator (PSR-36, Neuralynx Inc.) above the arena, allowing free turning movement of the animal. From there, the signals were relayed to a second amplifier in the adjacent room. Signals were amplified by a factor of 3000—5000, band-pass filtered between 0.3 and 6 kHz and saved to computer for offline analysis. One red light emitting diode situated on the top of the headstage was used for tracking the rat’s head position and was monitored at 25 Hz by an overhead camera and a digital spot follower. The electrodes were not moved after the end of the recordings.

### Data Analyses

After acquisition, all 32 channels were visually screened to identify putative units and thresholds were set in order to digitize spike waveforms at 32 kHz for 1 ms. In this way, even low firing units were successfully isolated for every recording session. If multiple sessions were ran successively, the same thresholds were applied to all recordings to ensure correct identification of the same units over time. Clusters from different units were further refined using offline sorting tools (Offline Sorter^TM^, Plexon^®^). The quality of unit isolation was assessed by using several methods, as follows: first, autocorrelation histograms were calculated for each unit, and the unit was removed from further analysis if the histogram revealed the existence of correlations within the first 2 ms (refractory period), inconsistent with good unit isolation. Second, for cells recorded simultaneously on the same electrode, we measured the degree to which the selected unit clusters are separated in the multidimensional cluster views as determined by a Multivariate Analysis of Variance (MANOVA) test (Offline Sorter^TM^). Sorted clusters were kept for analysis only if MANOVA tests yielded a *p* value < 0.001.

### Spatial Firing Analysis

Firing rate maps allow for visual inspection of neurons preferred areas of firing (*i.e.* place fields). They were constructed by dividing the number of spikes which occurred in specific pixel coordinates by the total trial time the animal spent in that coordinate. This produced maps depicting the place fields of each cell in Hertz. The pixel map is converted into a 30×30 array of square bins 2.5 cm on a side. Autoscaled color-coded firing rate maps were then created to visualize firing rate distributions^1^. In such maps, pixels in which no spikes occurred during the whole session are displayed as blue. The highest firing rate is coded as red, and intermediate rates are shown as orange, yellow, green, and cyan pixels from high to low. We used multiple indices to analyse the spatial properties of the hippocampus place cell firing (namely spatial coherence and spatial information content). A firing field was defined as a set of at least nine contiguous pixels with firing rate above zero. Only the largest field is considered for each cell if more than one field was found. The centre of the field was defined as the 3×3 group of pixels with the greatest mean rate^2^. Spatial coherence consists of a spatial autocorrelation of the place field map and measures the extent to which the firing rate in a particular bin is predicted by the average rate of the eight surrounding bins. Thus, high positive values resulted if the rate for each bin could be better predicted given the firing frequency of the neighbour location^3^. The spatial information content is expressed in bits per spike^4^ and is calculated as follows:

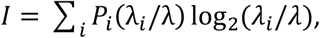

where *λ*_*i*_ is the mean firing rate in bin *i*, *λ* is the overall mean firing rate and *P*_*i*_ is the occupancy probability of bin *i*. In short, the spatial information content index can be seen as a measure of the amount of information relative to the location of the animal conveyed by a single action potential emitted by a single place cell. Similar to others^5,6^, place cells were selected for study if their spatial firing patterns were location-specific (coherence ≥ 0.5; spatial information ≥ 0.5 bits/AP) and robust (average activity ≥ 0.25Hz).

In order to assess the remapping events, we performed cross-correlations between the smoothed firing rate maps (using a 5×5 kernel) of the sessions where the same neurons have been identified based on their waveform properties^7^. To be considered as a remapping event the correlation had to fall under a cut-off of *r* = 0.5 ^7,8^.

### Overdispersion

Overdispersion was measured as described in Fenton and Muller^9^. Passes through the firing field were defined as the time series of positions starting when the light-emitting diode (LED) was detected inside the field and ending when the LED was detected outside the field. To enhance the reliability of firing rate estimates, passes were studied only if they met the two following criteria: (*i*) each pass had to last at least 1 second; (*ii*) the pass had to go through the field center. The observed number of spikes fired during a pass was compared with the number of spikes predicted from the session-averaged positional firing rate distribution. The predicted activity during a pass depends only on the specific pixels visited and the time spent in those pixels without regard to the sequence of positions.

For a given pass, the expected number of spikes is given by:

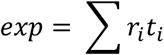

where *r*_*i*_ is the time-averaged firing rate at position in the pass through the field, and *t*_*i*_ the time spent in location *i* during the pass. According to the Poisson assumption, the standard deviation of the expected numbers of spikes is equal to 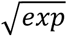. Thus *Z* is a standard-normal deviate that measures the standardized deviation of the observed discharge (*obs*) from this expectation for each pass, and is calculated as follows:

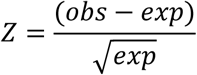

Therefore, if *Z* ≥ 1.96 or ≤ 1.96, the probability that the observed number of spikes is consistent with the model is inferior or equal to 0.05. Overdispersion was then measured as the variance of the distribution of *Z* values computed for a set of passes.

### Local field potentials analysis

Local field potentials (LFP) were recorded between two electrodes along with single units, lowpass-filtered at 475 Hz and sampled at 32 kHz. In order to characterize the theta oscillations during exploration of the arena, we resampled the signal to 256Hz, applied a band-pass filter with cut-off frequencies in the theta band (4-12 Hz) and in the delta band (2-4 Hz; FMA Toolbox distributed under General Public License, http://fmatoolbox.sourceforge.net). We computed the Hilbert transform of the resulting oscillations and we calculated the instantaneous frequency from which we extracted the peak frequency in theta and delta bands. In order to compare the relative power of theta oscillations between CA3 and dLGN recordings, we calculated the theta/(theta+delta) ratio by dividing the theta peak power by the sum of theta and delta peak powers. Finally, intrinsic firing frequency of place cells in the theta band (4-12 Hz) was calculated by extracting the theta peak of the Hilbert transform of the cell spike autocorrelogram.

### Phase precession

Two-dimensional phase precession of place cells was computed in the arena using the ‘pass-index’ Matlab function distributed under General Public License (https://github.com/jrclimer/Pass_Index) and developed by Climer et al.^10^. To compute the field index we first calculated occupancy-normalized firing rate for 1cm square bins of position data. Data were then smoothed with a pseudo-Gaussian kernel with a five pixel (5 cm) standard deviation. Each bin of the field index map was then attributed a value percentile normalized between 0 and 1. The omnidirectional pass index was computed on the raw rate maps. Quantification of phase precession was achieved by using the circular-linear correlation coefficient^11^ (*rho*), the statistical significance of the Pearson correlation (*p* value) and the slope (*s* expressed in degrees per pass). A place cell was considered omnidirectionally precessing if the significance level of the correlation was below 0.01.

### 1 Hz power modulation

Intrinsic firing frequency of cells was extracted from the Fast Fourier transform (FFT) of the cell spike-time autocorrelation sequence with 50ms bins. Cells with a 1Hz power value greater than five times the average of the power of the whole FFT band was classified as a 1 Hz modulated cell.

### Spatial sparsity

In order to compare spatial selectivity between place cells and 1Hz modulated cells, we employed an index independent of the criteria used to identify place cells. Such index, the sparsity of a rate map, was defined as follows^12^:

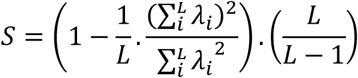

where *λ*_*i*_ is the firing rate in the *i*-th bin, and *L* is the number of spatial bins of the rate map. Thus defined, *S* ranges between zero (for a uniform rate map) and 1 (for a delta function rate map).

### Histology

At the end of the experiments, rats were killed by an injection of an overdose of sodium pentobarbital (180 mg/kg, i.p.), then transcardially perfused with a 4% phosphate-buffered (0.1 m) paraformaldehyde solution at 4°C. Their brains were removed, postfixed in the same fixative at 4°C during 2 h, and transferred to a 30% phosphate-buffered (0.1 m) sucrose solution for 48 h at 4°C before being snap frozen using isopentane at −40°C. They were then stored at −80°C before further processing. Using a cryostat, coronal sections (40 μm) were cut within a block of tissue extending from −3.00 to −4.92 mm from bregma^13^ to allow the evaluation of both CA3 and dLGN tetrode implantations. All of the sections were collected on gelatin-coated slides and processed for cresyl violet staining. The positions of the tips of the electrodes were determined from digital pictures, acquired with a Leica Microscope (Wetzlar, Germany) and imported in an image manipulation program (Gimp 2.8, distributed under General Public License). The depth of the different recordings was reconstructed based on the deepest positions of the electrode traces and daily recorded information of electrode position.

### Statistics

No statistical methods were used to predetermine sample sizes, but our sample sizes are similar to those reported for place cells in previous studies^7,14^. Statistical tests included Levene’s test for equality of variances, two-sample t-test assuming unequal variances, two-sample t-test assuming equal variances, Pearson’s correlations and χ^2^ test. Data distribution was assumed to be normal, but this was not formally tested. A Supplementary Methods Checklist is available. Circular statistics were based on Rayleigh tests and circular V-tests. To measure whether cell activity was controlled by apparatus landmarks between two recording sessions, a cross-correlation procedure was used by rotating one ratemap in steps of 10° and calculating the Pearson correlation coefficient between this rotated and the second unrotated rate map at each steps. The correlation value at 0° step was used as a measure of spatial stability or remapping. The angle with the greater correlation coefficient was also extracted. For sessions where the set of objects was rotated by an amount of 90°, the data was analyzed by using a circular V-test with 90° as a specified direction. When no rotation was made, the same test was performed with 0° as a specified direction. The Rayleigh vector length and the circular mean direction were calculating using circular statistic toolbox^15^ (Matlab, CircStat: A MATLAB Toolbox for Circular Statistics distributed under General Public License, https://www.jstatsoft.org/article/view/v031i10).

## Supplementary Discussion

The discovery of spatially tuned neurons with complementary functions in the brain (i.e. hippocampal place cells^1–3^, head direction cells^4,5^ and grid cells^6,7^) suggests the existence of a vast network specialized in the processing of spatial information. Surprisingly, given the central role of visual information in orientation behaviors, few studies so far explored the direct role of visual areas in spatial cognition. For instance, seminal work by Tsang^8^ demonstrated that early blind rats trained adults in a maze and subsequently lesioned in the visual cortex showed profound disruption in their maze ability. Subsequent studies by Lashley^9^ further supported the evidence that the primary visual area (V1) was necessary to form a spatial representation of some kind and that it is used in the learning process. More recently, Goodale and Dale^10^ showed that visually brain-damaged rats were much more impaired in a spatial task than blind rats, while Paz-Villagrán et al.^11^ showed that lesions of the primary visual area provoked a strong internal disorganization of hippocampal place fields. Investigating the role of the primary visual area in spatial cognition is further motivated by recent studies showing a clear spatial selectivity of visual cortical neurons in freely moving rodents^12,13^. More importantly, these cortical neurons in V1 show also robust replay events^13^.

Aside from the hippocampo-cortical interactions, there is growing evidence that the thalamus plays a key role in dynamically routing information across the brain^14,15^. Such a role may involve flexibly synchronizing ensembles of neurons, thereby configuring brain networks for the current behavioral context^16^. It appears therefore that thalamic interactions with cortical and subcortical structures are involved in various processes such as behavioral flexibility, memory formation and cognition in general.

The dorsolateral geniculate nucleus (dLGN – thalamic structure located one synapse upstream of the primary visual area) is classically considered as the main thalamic gateway for visual information from the retina to reach the cerebral cortex^17^. However, over 90% of rat retinal ganglion cells project to the superior colliculus (SC), whereas a smaller proportion (about 20%) project to the dLGN and thus to V1^18,19^. As pointed out by A. David Milner in his open peer commentary of Foreman and Stevens article^20^, it is not unreasonable to think of the superior colliculus as processing and providing inputs necessary for higher-level spatial representations. In the main, SC transforms retinotopic coordinates of particular events in egocentric (body-centered) space in order to direct behavioral responses toward these specific events. Interestingly, it has also been shown that the outer shell of the dLGN receives a strong input from SC^21^. Other brain stem structures appear to project onto the dLGN, and more specifically the pedunculopontine tegmental (PPT) nucleus^22^ known to be involved, among other things, in the modulation of locomotor behavior^23^. Therefore, dLGN has virtually access to all components necessary to build up an allocentric representation of space^24^, that is instrumental in creating a global navigation system.

Following this idea, we asked in the present study whether cells in the dLGN would show spatial selectivity while the animals are foraging in a circular arena. Indeed, we found numerous place cells in the dLGN showing coding properties thought until then hallmarks of hippocampal function (*e.g.*, global remapping). Moreover, the spatial selectivity of dLGN cells was maintained in the absence of visual inputs, i.e. when rats were placed in complete darkness, thus demonstrating that these spatially localized firing patterns, once learned, are not likely to depend upon local visual cues within the experimental room. In a parallel experiment, we also found non-spatial dLGN cells responding to full-field visual stimulations (ambient light flickering at 1Hz) recorded simultaneously with place cells, thus providing a complementary hint that these cells were indeed recorded in the dLGN. It appears therefore that dLGN might play a much broader role in spatial cognition than mere visual processing brain structure.

Based on our results, we identified several key questions that will have to be addressed in the near future. Given the strong feedback connections originating from V1^25^, one has to determine if the dLGN spatial selectivity that we observed is dependent upon visual cortex integrity. Reversible silencing appears therefore as a powerful approach for this purpose. Optogenetic manipulations can achieve this by rapidly modulating the activity of specific brain regions and cell types. Indeed, infecting and activating PV+ interneurons with channelrhodopsin-2 has been shown to provide effective transient suppression of neuronal activity in primary visual cortex^26^.

Another key question concerns the ability of dLGN to participate to stabilization of a previously encoded memory trace. Hippocampal replay has been first observed during rapid eye movement sleep periods^27^ that followed initial exposure to an environment. This phenomenon has been also described during wakefulness^28^ and more specifically during an EEG state characterized by sharp waves and high-frequency oscillations (ripples). During replay, place cells that fired during exposure to an environment are orderly reactivated at a subsequent time so that the initial experience is recapitulated over a very brief period of time, a phenomenon taken to reflect the operation of an offline consolidation mechanism^29^. It appears clearly now that the hippocampus is not the only structure showing such features, as previous work showed that neurons in V1 possessed a replay activity during sleep^13^. Following Buzsáki and colleagues hypothesis^30^, the presence of replay within the hippocampus could serve to perform “what if” scenarios and anticipate the possible consequences of alternative actions without actually testing them. If replay is also observed in dLGN, this would mean that this structure, a key component in visual processing, would indeed perform computations largely independent of pure sensory inputs. This would however raise intriguing questions concerning the functional differentiation between hippocampus and dLGN. Our current view is that the dLGN acts as the true mapping structure of the navigation system, representing space as a passive computation, while the hippocampus supports an active spatial process involved in route planning^31,32^.

Given the primary role in visual processing of the dLGN, a fine characterization of the visual receptive field properties of these neurons is primordial. Their properties are indeed very well described in the literature, with specific center-surround antagonistic receptive field profiles^33,34^, that are only shared by retinal ganglion cells in the visual system. The central question, here, would be to ask whether these neurons share any particular visual properties (e.g. like magnocellular thalamic neurons with large receptive field, sensitivity to motion, as opposed to parvocellular neurons) that could be functionally related to their sensitivity to place.

Lastly, we feel that this work and subsequent studies in the field will be directly relevant to a growing list of disorders, in particular neurodegenerative diseases, for which dysfunctional inter-structure communication and plasticity may be a major cause of cognitive deficits. For instance, visual hallucination is a common non-motor symptom (ca. 20% of occurrence) in patients with Parkinson’s disease^35^. Prevalence of visual hallucination increases as Parkinson’s disease progresses^36^, and the presence of this ailment is predictive of poor outcomes, such as those related to cognitive impairment, institutionalization, and higher mortality^37^. Recent works highlight the crucial role of the lateral geniculate nucleus as patients with visual hallucinations showed a clear atrophy at this level^38^. In the main, visual hallucinations are thought to be linked to a lack of suppression or spontaneous emergence of internally generated imagery through the ponto-geniculo-occipital system^39^. Investigating such network through the spatial cognition paradigm as we suggest might help to understand the etiology of visuospatial cognitive deficits that accompany sometimes visual hallucinations^40,41^. Overall, the network relying on brain structures classically defined as visual structures (e.g. primary visual cortex, dorsal lateral geniculate nucleus and superior colliculus) may also be a prime candidate for future studies in the primate field of research on memory.

**Author contributions**
B.P., E.S. and V.H. planned and interpreted the study, P.B., B.T. and V.H. performed surgeries and acquired the data. P-Y.J. and V.H. carried out the analyses. P-Y.J, B.T., B.P., E.S. and V.H. wrote the paper.

## Acknowledgements

Support for this work was provided by the Centre National de la Recherche Scientifique (CNRS) and the Agence Nationale de la Recherche (ANR-10-BLAN-1413). The authors wish to thank Frédéric Chavane and Matteo Carandini for useful discussions, Francesca Sargolini for her careful reading and suggestions, Laure Spieser for her help in programming and Didier Louber for his precious technical assistance. P-Y.J. salary was financed by the Fondation pour la Recherche Médicale (N° ARF20170938640).

## References

1. O’Keefe, J. & Dostrovsky, J. The hippocampus as a spatial map. Preliminary evidence from unit activity in the freely-moving rat. Brain Res. 34, 171–175 (1971).

2. Ekstrom, A. D. et al. Cellular networks underlying human spatial navigation. Nature 425, 184–8 (2003).

3. Epstein, R. A., Patai, E. Z., Julian, J. B. & Spiers, H. J. The cognitive map in humans: spatial navigation and beyond. Nat. Neurosci. 20, 1504–1513 (2017).

4. Moser, E. I., Kropff, E. & Moser, M.-B. Place cells, grid cells, and the brain’s spatial representation system. Annu Rev Neurosci 31, 69–89 (2008).

5. Moser, E. I., Moser, M.-B. & McNaughton, B. L. Spatial representation in the hippocampal formation: a history. Nat. Neurosci. 20, 1448–1464 (2017).

6. Jankowski, M. M. & O’Mara, S. M. Dynamics of place, boundary and object encoding in rat anterior claustrum. Front. Behav. Neurosci. 9, 250 (2015).

7. Mao, D., Kandler, S., McNaughton, B. L. & Bonin, V. Sparse orthogonal population representation of spatial context in the retrosplenial cortex. Nat. Commun. 8, 243 (2017).

8. Ji, D. & Wilson, M. A. Coordinated memory replay in the visual cortex and hippocampus during sleep. Nat Neurosci 10, 100–107 (2007).

9. Haggerty, D. C. & Ji, D. Activities of visual cortical and hippocampal neurons co-fluctuate in freely moving rats during spatial navigation. eLife e08902 (2015). doi:10.7554/eLife.08902

10. Saleem, A. B., Diamanti, E. M., Fournier, J., Harris, K. D. & Carandini, M. Coherent encoding of subjective spatial position in visual cortex and hippocampus. Nature (2018). doi:10.1038/s41586-018-0516-1

11. Barry, J. M. Axonal activity in vivo: technical considerations and implications for the exploration of neural circuits in freely moving animals. Front. Neurosci. 9, 153 (2015).

12. Tsao, A., Moser, M.-B. & Moser, E. I. Traces of experience in the lateral entorhinal cortex. Curr. Biol. CB 23, 399–405 (2013).

13. Hok, V., Chah, E., Reilly, R. B. & O’Mara, S. M. Hippocampal dynamics predict interindividual cognitive differences in rats. J Neurosci 32, 3540–3551 (2012).

14. Fenton, A. A. et al. Attention-like modulation of hippocampus place cell discharge. J Neurosci 30, 4613–4625 (2010).

15. Jackson, J. & Redish, A. D. Network dynamics of hippocampal cell-assemblies resemble multiple spatial maps within single tasks. Hippocampus 17, 1209–1229 (2007).

16. Ravassard, P. et al. Multisensory Control of Hippocampal Spatiotemporal Selectivity. Science 340, 1342–1346 (2013).

17. Muller, R. U. & Kubie, J. L. The effects of changes in the environment on the spatial firing of hippocampal complex-spike cells. J Neurosci 7, 1951–68 (1987).

18. Leutgeb, J. K., Leutgeb, S., Moser, M.-B. & Moser, E. I. Pattern separation in the dentate gyrus and CA3 of the hippocampus. Science 315, 961–966 (2007).

19. Colgin, L. L., Moser, E. I. & Moser, M.-B. Understanding memory through hippocampal remapping. Trends Neurosci 31, 469–477 (2008).

20. Killian, N. J., Jutras, M. J. & Buffalo, E. A. A map of visual space in the primate entorhinal cortex. Nature 491, 761–764 (2012).

21. Wilming, N., König, P., König, S. & Buffalo, E. A. Entorhinal cortex receptive fields are modulated by spatial attention, even without movement. eLife 7, (2018).

22. Haas, O. V., Henke, J., Leibold, C. & Thurley, K. Modality-specific Subpopulations of Place Fields Coexist in the Hippocampus. Cereb. Cortex (2018). doi:10.1093/cercor/bhy017

23. Monavarfeshani, A., Sabbagh, U. & Fox, M. A. Not a one-trick pony: Diverse connectivity and functions of the rodent lateral geniculate complex. Vis. Neurosci. 34, E012 (2017).

24. Zhao, X., Chen, H., Liu, X. & Cang, J. Orientation-selective responses in the mouse lateral geniculate nucleus. J Neurosci 33, 12751–12763 (2013).

25. Olsen, S. R., Bortone, D. S., Adesnik, H. & Scanziani, M. Gain control by layer six in cortical circuits of vision. Nature 483, 47–52 (2012).

26. Aguila, J., Cudeiro, F. J. & Rivadulla, C. Suppression of V1 Feedback Produces a Shift in the Topographic Representation of Receptive Fields of LGN Cells by Unmasking Latent Retinal Drives. Cereb. Cortex 27, 3331–3345 (2017).

27. Kerschensteiner, D. & Guido, W. Organization of the dorsal lateral geniculate nucleus in the mouse. Vis. Neurosci. 34, (2017).

28. Wilson, J. J., Alexandre, N., Trentin, C. & Tripodi, M. Three-Dimensional Representation of Motor Space in the Mouse Superior Colliculus. Curr. Biol. CB 28, 1744–1755.e12 (2018).

29. Erişir, A., Van Horn, S. C. & Sherman, S. M. Relative numbers of cortical and brainstem inputs to the lateral geniculate nucleus. Proc. Natl. Acad. Sci. U. S. A. 94, 1517–1520 (1997).

30. Cressant, A., Muller, R. U. & Poucet, B. Failure of centrally placed objects to control the firing fields of hippocampal place cells. J Neurosci 17, 2531–42 (1997).

## Supplementary Methods References

1. Muller, R. U., Kubie, J. L. & Ranck, J. B. Spatial firing patterns of hippocampal complex-spike cells in a fixed environment. J Neurosci 7, 1935–50 (1987).

2. Fenton, A. A., Csizmadia, G. & Muller, R. U. Conjoint control of hippocampal place cell firing by two visual stimuli. I. The effects of moving the stimuli on firing field positions. J Gen Physiol 116, 191–209 (2000).

3. Muller, R. U. & Kubie, J. L. The firing of hippocampal place cells predicts the future position of freely moving rats. J Neurosci 9, 4101–10 (1989).

4. Skaggs, W. E., McNaughton, B. L. & Gothard, K. M. An Information-Theoretic Approach to Deciphering the Hippocampal Code. in (eds. Hanson, S. J., Cowan, J. D. & Giles, C. L.) 5, 1030–1037 (Morgan Kaufmann, San Mateo, CA, 1993).

5. Leutgeb, J. K., Leutgeb, S., Moser, M.-B. & Moser, E. I. Pattern separation in the dentate gyrus and CA3 of the hippocampus. Science 315, 961–966 (2007).

6. Fenton, A. A. et al. Attention-like modulation of hippocampus place cell discharge. J Neurosci 30, 4613–4625 (2010).

7. Hok, V., Chah, E., Reilly, R. B. & O’Mara, S. M. Hippocampal dynamics predict interindividual cognitive differences in rats. J Neurosci 32, 3540–3551 (2012).

8. Barnes, C. A., Suster, M. S., Shen, J. & McNaughton, B. L. Multistability of cognitive maps in the hippocampus of old rats. Nature 388, 272–275 (1997).

9. Fenton, A. A. & Muller, R. U. Place cell discharge is extremely variable during individual passes of the rat through the firing field. Proc Natl Acad Sci U A 95, 3182–7 (1998).

10. Climer, J. R., Newman, E. L. & Hasselmo, M. E. Phase coding by grid cells in unconstrained environments: two-dimensional phase precession. Eur. J. Neurosci. 38, 2526–2541 (2013).

11. Kempter, R., Leibold, C., Buzsáki, G., Diba, K. & Schmidt, R. Quantifying circular–linear associations: Hippocampal phase precession. J. Neurosci. Methods 207, 113–124 (2012).

12. Ravassard, P. et al. Multisensory Control of Hippocampal Spatiotemporal Selectivity. Science 340, 1342–1346 (2013).

13. Paxinos, G. & Watson, C. The rat brain in stereotaxic coordinates. (Elsevier, Amsterdam:, 2007).

14. Hok, V. et al. Goal-related activity in hippocampal place cells. J Neurosci 27, 472–482 (2007).

15. Berens, P. CircStat : A MATLAB Toolbox for Circular Statistics. J. Stat. Softw. 31, (2009).

## Supplementary Discussion References

3. Muller, R. U., Kubie, J. L. & Ranck, J. B. Spatial firing patterns of hippocampal complex-spike cells in a fixed environment. J Neurosci 7, 1935–50 (1987).

4. Ranck, J. B. Head direction cells in the deep cell layer of dorsal presubiculum in freely moving rats. Soc Neurosci Abstr 10, 599–599 (1984).

5. Taube, J.S. The Head Direction Signal: Origins and Sensory-Motor Integration. Annu Rev Neurosci 30, 181–207 (2007).

6. Hafting, T., Fyhn, M., Molden, S., Moser, M.-B. & Moser, E. I. Microstructure of a spatial map in the entorhinal cortex. Nature 436, 801–806 (2005).

7. Boccara, C. N. et al. Grid cells in pre- and parasubiculum. Nat Neurosci 13, 987–994 (2010).

8. Tsang, Y.-C. The functions of the visual areas of the cerebral cortex of the rat in the learning and retention of the maze. I. I. (private edition : distributed by The University of Chicago libraries, 1934).

9. Lashley, K. S. Studies of cerebral function in learning XII. Loss of the maze habit after occipital lesions in blind rats. J. Comp. Neurol. 79, 431–462 (1943).

10. Goodale, M. A. & Dale, R. H. Radial-maze performance in the rat following lesions of posterior neocortex. Behav. Brain Res. 3, 273–288 (1981).

11. Paz-Villagrán, V., Lenck-Santini, P. P., Save, E. & Poucet, B. Properties of place cell firing after damage to the visual cortex. Eur J Neurosci 16, 771–6 (2002).

12. Haggerty, D. C. & Ji, D. Activities of visual cortical and hippocampal neurons co-fluctuate in freely moving rats during spatial navigation. eLife e08902 (2015). doi:10.7554/eLife.08902

13. Ji, D. & Wilson, M. A. Coordinated memory replay in the visual cortex and hippocampus during sleep. Nat Neurosci 10, 100–107 (2007).

14. Loureiro, M. et al. The Ventral Midline Thalamus (Reuniens and Rhomboid Nuclei) Contributes to the Persistence of Spatial Memory in Rats. J. Neurosci. 32, 9947–9959 (2012).

15. Xu, W. & Südhof, T. C. A Neural Circuit for Memory Specificity and Generalization. Science 339, 1290–1295 (2013).

16. Saalmann, Y. B. & Kastner, S. The cognitive thalamus. Front. Syst. Neurosci. 9, (2015).

17. Groenewegen, H. J. & Witter, M. P. Chapter 17 - Thalamus A2 - Paxinos, George. in The Rat Nervous System (Third Edition) 407–453 (Academic Press, 2004).

18. Dreher, B., Sefton, A. J., Ni, S. Y. & Nisbett, G. The morphology, number, distribution and central projections of Class I retinal ganglion cells in albino and hooded rats. Brain. Behav. Evol. 26, 10–48 (1985).

19. Linden, R. & Perry, V. H. Massive retinotectal projection in rats. Brain Res. 272, 145–149 (1983).

20. Foreman, N. & Stevens, R. Relationships between the superior colliculus and hippocampus: Neural and behavioral considerations. Behav. Brain Sci. 10, 101 (1987).

21. Reese, B. E. The projection from the superior colliculus to the dorsal lateral geniculate nucleus in the rat. Brain Res. 305, 162–168 (1984).

22. Rye, D. B., Saper, C. B., Lee, H. J. & Wainer, B. H. Pedunculopontine tegmental nucleus of the rat: cytoarchitecture, cytochemistry, and some extrapyramidal connections of the mesopontine tegmentum. J. Comp. Neurol. 259, 483–528 (1987).

23. Garcia-Rill, E., Houser, C. R., Skinner, R. D., Smith, W. & Woodward, D. J. Locomotion-inducing sites in the vicinity of the pedunculopontine nucleus. Brain Res. Bull. 18, 731–738 (1987).

24. Poucet, B. Spatial cognitive maps in animals: new hypotheses on their structure and neural mechanisms. Psychol Rev 100, 163–82 (1993).

25. Bourassa, J. & Deschênes, M. Corticothalamic projections from the primary visual cortex in rats: a single fiber study using biocytin as an anterograde tracer. Neuroscience 66, 253–263 (1995).

26. Glickfeld, L. L., Histed, M. H. & Maunsell, J. H. R. Mouse primary visual cortex is used to detect both orientation and contrast changes. J Neurosci 33, 19416–19422 (2013).

27. Louie, K. & Wilson, M. A. Temporally Structured Replay of Awake Hippocampal Ensemble Activity during Rapid Eye Movement Sleep. Neuron 29, 145–156 (2001).

28. Foster, D. J. & Wilson, M. A. Reverse replay of behavioural sequences in hippocampal place cells during the awake state. Nature 440, 680–683 (2006).

29. Girardeau, G., Benchenane, K., Wiener, S. I., Buzsáki, G. & Zugaro, M. B. Selective suppression of hippocampal ripples impairs spatial memory. Nat. Neurosci. 12, 1222–1223 (2009).

30. Buzsáki, G., Peyrache, A. & Kubie, J. Emergence of Cognition from Action. Cold Spring Harb. Symp. Quant. Biol. 79, 41–50 (2014).

31. Grieves, R. M., Wood, E. R. & Dudchenko, P. A. Place cells on a maze encode routes rather than destinations. eLife 5, (2016).

32. Spiers, H. J., Olafsdottir, H. F. & Lever, C. Hippocampal CA1 activity correlated with the distance to the goal and navigation performance. Hippocampus (2017). doi:10.1002/hipo.22813

33. Grieve, K. L. Binocular visual responses in cells of the rat dLGN. J. Physiol. 566, 119–124 (2005).

34. Sriram, B., Meier, P. M. & Reinagel, P. Temporal and spatial tuning of dorsal lateral geniculate nucleus neurons in unanesthetized rats. J. Neurophysiol. 115, 2658–2671 (2016).

35. Khoo, T. K. et al. The spectrum of nonmotor symptoms in early Parkinson disease. Neurology 80, 276–281 (2013).

36. Fénelon, G., Mahieux, F., Huon, R. & Ziégler, M. Hallucinations in Parkinson’s disease: prevalence, phenomenology and risk factors. Brain J. Neurol. 123 (Pt 4), 733–745 (2000).

37. Hely, M. A., Reid, W. G. J., Adena, M. A., Halliday, G. M. & Morris, J. G. L. The Sydney multicenter study of Parkinson’s disease: the inevitability of dementia at 20 years. Mov. Disord. Off. J. Mov. Disord. Soc. 23, 837–844 (2008).

38. Lee, J.-Y. et al. Lateral geniculate atrophy in Parkinson’s with visual hallucination: A trans-synaptic degeneration? Mov. Disord. Off. J. Mov. Disord. Soc. 31, 547–554 (2016).

39. Diederich, N. J., Goetz, C. G. & Stebbins, G. T. Repeated visual hallucinations in Parkinson’s disease as disturbed external/internal perceptions: focused review and a new integrative model. Mov. Disord. Off. J. Mov. Disord. Soc. 20, 130–140 (2005).

40. Davidsdottir, S., Cronin-Golomb, A. & Lee, A. Visual and spatial symptoms in Parkinson’s disease. Vision Res. 45, 1285–1296 (2005).

41. Ozer, F. et al. Cognitive impairment patterns in Parkinson’s disease with visual hallucinations. J. Clin. Neurosci. Off. J. Neurosurg. Soc. Australas. 14, 742–746 (2007).

